# Proficiency-Dependent Reorganisation of Language and Control Networks during Second Language Processing: An fMRI Study of Korean–English Bilinguals

**DOI:** 10.64898/2026.08.14.744902

**Authors:** Joonwoo Kim, Jiyoun Choi, Yeonji Baik, Walter J. B. van Heuven, Kichun Nam, JeYoung Jung

## Abstract

Second language (L2) processing engages both language-specific and domain-general control systems, yet how these systems vary with L2 proficiency remains unclear. We used functional magnetic resonance imaging (fMRI) to examine neural activity during L2 English processing in Korean–English (K-E) bilinguals across three proficiency levels (beginner, intermediate, advanced). Participants performed rhyme and spelling judgement tasks manipulating orthographic–phonological conflict. Behaviourally, conflict conditions reduced accuracy, with proficiency effects observed selectively in the rhyme task. fMRI results showed that conflict processing recruited frontoparietal control regions, including inferior frontal and parietal cortices, accompanied by deactivation in default mode network regions. Critically, proficiency-related effects differed by task. During rhyme judgement, advanced bilinguals showed greater activation in the left supramarginal gyrus (SMG) and cerebellum, whereas intermediate bilinguals exhibited greater recruitment of the left middle orbital gyrus and dorsomedial prefrontal cortex. During spelling judgement, advanced bilinguals showed greater thalamic activation alongside greater deactivation of the right dorsolateral prefrontal cortex. Activity in the left SMG and cerebellum was positively associated with L2 reading score, and cerebellar activity was also associated with rhyme-task performance, whereas right DLPFC activity was negatively associated with the scores. These findings suggest that increasing L2 proficiency is associated less with altered recruitment of core reading regions than with task-specific shifts in the balance between phonological-specialized, subcortical attentional, and domain-general control systems supporting L2 processing.

## 1. Introduction

Bilingualism is a widespread phenomenon worldwide, in which individuals acquire and use more than one language (Grosjean, 1998). A central question in the cognitive neuroscience of bilingualism concerns how the human brain accommodates a later-acquired second language (L2), and how the neural systems supporting L2 processing vary across individuals (Abutalebi et al., 2001; Indefrey, 2006; Klein et al., 2014; Kroll and de Groot, 2005; Sebastian et al., 2011; Stowe and Sabourin, 2005; van Heuven and Dijkstra, 2010). Among the multiple factors that shape this variation, L2 proficiency has been identified as one of the most influential moderators of both behavioural performance and neural activity (Abutalebi, 2008; Luk and Bialystok, 2013; Mishra, 2014; Sulpizio et al., 2020). Even within the same language pair, bilinguals with different proficiency levels often show distinct patterns of cortical recruitment during L2 processing, suggesting that proficiency-related variation reflects systematic differences in neural engagement. Despite a broad consensus on its centrality, the neural basis of within-bilingual variation in L2 proficiency remains incompletely characterised, particularly across different linguistic domains and in less-studied bilingual populations (Takahesu Tabori et al., 2018).

Behavioural studies indicate that L2 proficiency varies across linguistic domains. Phonological processing is particularly sensitive to L2 proficiency, with lower-proficiency bilinguals showing slower and less accurate performance during tasks requiring phonological manipulation or resolution of orthographic and phonological conflict (e.g., *mint–pint*, *light–kite*) (Cao et al., 2013; Wang et al., 2009; Wartenburger et al., 2003). Orthographic processing appears more robust to proficiency differences at the behavioural level, but it remains sensitive to the typological distance between the first language (L1) and the L2, particularly when the two languages differ substantially in orthographic depth (Meschyan and Hernandez, 2006; Sulpizio et al., 2020; van Heuven and Dijkstra, 2010). Importantly, neuroimaging studies frequently reveal proficiency-related differences even when behavioural performance is comparable across groups with different levels of proficiency, suggesting that functional imaging may capture compensatory or efficiency-related mechanisms not evident behaviourally (Cargnelutti et al., 2019).

Functional neuroimaging studies have shown that L1 and L2 processing recruit a largely overlapping language network (Buchweitz et al., 2012; Chee et al., 1999; Indefrey, 2006; Klein et al., 1999; Mei et al., 2015; Tan et al., 2003), including the left inferior frontal gyrus (IFG), superior temporal gyrus (STG), supramarginal gyrus (SMG), inferior temporal gyrus (ITG), middle temporal gyrus (MTG), fusiform gyrus (FG), and superior parietal lobule (SPL) (Booth et al., 2004; Callan et al., 2004; Chen and Desmond, 2005; Jones et al., 2012; Price, 2010). However, L2 processing often elicits stronger or more widespread activation than L1, particularly for less-proficient or less-automatic L2 use (Cargnelutti et al., 2019; Luk and Bialystok, 2013; Perani et al., 2003, 1998). These differences extend beyond classical language regions to domain-general systems involved in cognitive control and attention.

In particular, frontoparietal and executive control systems are consistently engaged during L2 processing, reflecting increased demands on attentional regulation, conflict monitoring, and inhibition (Cargnelutti et al., 2019; Cole and Schneider, 2007; Grady et al., 2015). Subcortical structures such as the thalamus and basal ganglia have also been implicated in bilingual language control (Abutalebi and Green, 2007; Green, 1998), and interlingual conflict between a bilingual’s two languages has been shown to engage this control circuitry together with left prefrontal cortex (van Heuven et al., 2008). The default mode network (DMN), which typically deactivates during externally directed tasks, has been shown to deactivate more strongly during demanding L2 processing, consistent with the increased cognitive effort required (Andrews-Hanna, 2012; Spreng et al., 2013). Together, these findings suggest that variation in L2 proficiency may involve not only differential recruitment of classical language regions but also graded engagement of domain-general control and attentional systems.

Recent theoretical frameworks suggest that bilingual language experience is characterised by dynamic interactions between language-specific and domain-general systems. The Adaptive Control Hypothesis (ACH; Green and Abutalebi, 2013) proposes that bilingual language use recruits multiple control processes including goal maintenance, conflict monitoring, interference suppression, and salient cue detection, whose engagement varies according to language demands and interactional context. Complementing this account, the Dynamic Restructuring Model (DRM; DeLuca et al., 2019; Pliatsikas, 2020) proposes increasing L2 experience is associated with progressive neurobiological adaptation, beginning with cortical changes in language-related regions and gradually shifting toward greater involvement of subcortical systems and more efficient cortical-subcortical coupling. Similarly, the Bilingual Anterior to Posterior and Subcortical Shift (BAPSS) framework (Grundy et al., 2017) proposes that bilingual experience initially relies heavily on frontal executive systems before transitioning toward more posterior and subcortical recruitment as processing becomes increasingly automatic. Although these frameworks differ in granularity, they converge on the prediction that variation in L2 proficiency should be reflected not only within the core language network but also in broader reorganisation of control, attention, and subcortical systems.

Korean-English (K-E) bilinguals provide an informative test case for examining these processes as Korean and English differ substantially in linguistic and orthographic structure. Korean employs a highly transparent featural alphabetic syllabary, whereas English has a deep alphabetic orthography in which the phonological form of a word is derived both from probabilistic spelling–sound correspondences and, for words with inconsistent grapheme-to-phoneme mappings, through lexical mediation. This asymmetry implies that K-E bilinguals may need to override an L1-driven, transparent grapheme-phoneme strategy when reading English, especially when orthographic and phonological information conflict. Prior work has shown that neural organisation in K-E bilinguals is both sensitive to age-of-acquisition and proficiency (Kim et al., 1997). More recently, Jung et al. (2018) reported that L2 processing in K-E bilinguals engages not only canonical language regions but also a broader set of control-and attention-related areas, with patterns of engagement that vary with L2 proficiency. A complementary line of work in Chinese-English bilinguals (Wang et al., 2020) has further shown that L2 proficiency is positively correlated with brain activation in cognitive control areas during L2 picture naming, supporting the view that proficiency-related neural differences extend beyond the core language network. Together, these findings position K-E bilinguals as well suited for a systematic, within-group investigation of how L2 proficiency modulates the interaction between language-specific and domain-general neural systems, although such a comprehensive characterisation across the proficiency spectrum has not yet been undertaken.

In the present study, we used fMRI to examine how L2 proficiency modulates neural activity during L2 processing in K-E bilinguals at three proficiency levels (advanced, intermediate, beginner). Participants performed rhyme and spelling judgement tasks in L2 English, in which orthographic and phonological information were either congruent or incongruent at the level of the word pair (non-conflict vs. conflict; see Methods for the levels of conflict that this manipulation involves). The two tasks were chosen to dissociate phonological from orthographic demands, which prior work suggests may be differentially sensitive to L2 proficiency. Based on the literature reviewed above, we tested three predictions. First, in regions classically associated with reading, defined by prior meta-analyses (left IFG, SMA, IPS, IPL, STG, ITG, FG, IOG; Martin et al., 2015; Tan et al., 2005; see also Price, 2012), we expected differential activation as a function of L2 proficiency, which we evaluated via a region-of-interest (ROI) analysis. Second, beyond these classical reading regions, we expected proficiency-modulated engagement of regions belonging to the frontoparietal, attention network, and DMN, with the degree of activation varying systematically across the three proficiency groups. Third, we expected these proficiency-related differences in regional activation to be associated with bilinguals’ L2 reading performance, both task accuracy under conflict and standardised L2 reading scores, thereby linking the neural pattern to the behavioural phenotype of L2 proficiency.

## 2. Methods

### 2.1. Participants

A total of 54 native Korean L2 learners of English were recruited from Korea University, Seoul, Korea. Participants were assigned to groups on the basis of documented residence in an English-speaking country. The advanced group had lived or been schooled abroad for eight years or longer, the intermediate group for one to three years, and the beginner group had learned English solely through the Korean public education system, with no period of residence abroad and no more than one year of supplementary language instruction. The resulting groups were matched for age (*F*(*2*, 51) = 1.06, *p* = .353) and sex (χ^2^(2) = 0.420, Fisher’s exact *p* = .836): advanced (*N* = 17, 14 females, age 18–28 years, *M* = 21.6), intermediate (*N* = 20, 15 females, age 19–27, *M* = 21.5), and beginner group (*N* = 17, 14 females, age 19–27, *M* = 22.6) (**Table 1**).

**Table 1.** Means and standard deviations (SDs) of language background information and L2 proficiency scores for advanced, intermediate, and beginner groups.

| Measure | Advanced<br>(N=17) | Intermediate<br>(N=20) | Beginner<br>(N=17) | ANOVA results |
| --- | --- | --- | --- | --- |
| Age<br>(years) | 21.6<br>(2.6) | 21.5<br>(2.1) | 22.6<br>(2.9) | $F(2, 51) = 1.06, p = .353$ |
| Age of L2 acquisition<br>(years) | 5.6<br>(2.5) | 6.2<br>(2.1) | 8.6<br>(2.8) | $F(2, 51) = 7.56, p = .001^{b, c}$ |
| L2 immersion<br>(years) | 10.5<br>(2.4) | 1.8<br>(0.6) | 0.0<br>(0.0) | $F(2, 51) = 286.72, p < .001^{a, b, c}$ |
| TIWRE<br>(max = 50) | 44.3<br>(2.0) | 40.4<br>(2.8) | 36.6<br>(3.4) | $F(2, 51) = 32.47, p < .001^{a, b, c}$ |

*Note.* TIWRE = Test of Irregular Word Reading Efficiency. Superscripts indicate significant post-hoc pairwise comparisons:
<sup>a</sup> advanced vs. intermediate groups;
<sup>b</sup> intermediate vs. beginner groups;
<sup>c</sup> advanced vs. beginner groups.

Participants completed a language background questionnaire covering the age of acquisition (AoA) of English and the number of years spent immersed in an English-speaking environment. One-way ANOVAs confirmed the intended separation in L2 immersion (*F*(*2*, 51) = 286.72, *p* < .001), and additionally revealed a group difference in AoA (*F*(*2*, 51) = 7.56, *p* = .001). Post-hoc pairwise comparisons showed that AoA was significantly earlier for advanced (*M* = 5.6, *t*(32) = 3.34, *p* = .006) and intermediate groups (*M* = 6.2, *t*(35) = 3.15, *p* = .010) than for beginner group (*M* = 8.6), whereas there was no difference between advanced and intermediate groups (*t*(35) = 0.68, *p* = 1.000). Post-hoc pairwise comparisons further revealed that the mean duration of L2 immersion was longest in advanced group (*M* = 10.5), followed by intermediate group (*M* = 1.8), with beginner group having no experience (advanced vs. intermediate, *t*(35) = 15.88, *p* < .001; advanced vs. beginner, *t*(32) = 18.23, *p* < .001; intermediate vs. beginner, *t*(35) = 12.53, *p* < .001).

L2 English proficiency was assessed using the Test of Irregular Word Reading Efficiency (TIWRE), which assesses the ability to read phonetically irregular English words (Reynolds and Kamphaus, 2007). In the test, participants were presented with 39 irregular English words on paper and asked to read each word aloud. Test scores were calculated as the total number of correctly pronounced words. Following the test manual for adults, 11 points were added to each participant’s raw score; thus, the highest possible score was 50. A one-way ANOVA showed a significant group difference in TIWRE scores (*F*(2, 51) = 32.47, *p* < .001). Post-hoc comparisons revealed that advanced group obtained the highest scores (M = 44.3), followed by intermediate (*M* = 40.4), and beginner group (*M* = 36.6), with all pairwise differences reaching significance (advanced vs. intermediate, *t*(35) = 4.75, *p* < .001; advanced vs. beginner, *t*(32) = 8.11, *p* < .001; intermediate vs. beginner, *t*(35) = 3.76, *p* = .002).

Of the 54 participants, nine were excluded from the fMRI analyses due to head motion exceeding 2 mm translation and/or 2° rotation, leaving 45 participants with usable imaging data in both tasks (advanced *N* = 16, intermediate *N* = 14, beginner *N* = 15). The proportion of excluded participants did not differ significantly across proficiency groups (advanced 1/17, intermediate 6/20, beginner 2/17; Fisher– Freeman–Halton exact *p* = .156), and the excluded participants did not differ from the retained sample in age (*t*(52) = 1.25, *p* = .215), TIWRE score (*t*(52) = 0.99, *p* = .327), AoA (*t*(52) = 0.81, *p* = .424), or L2 immersion (*t*(52) = 1.02, *p* = .314), indicating that motion-related data loss was not systematically related to L2 proficiency. In the retained imaging subsample, the three groups remained matched for age (*F*(2, 42) = 1.64, *p* = .207) and the group difference in TIWRE score remained significant (*F*(2, 42) = 29.14, *p* < .001; advanced *M* = 44.1, intermediate *M* = 39.6, beginner *M* = 36.5, all pairwise comparisons *p* < .05, Bonferroni corrected). All behavioural analyses reported below are based on the full sample of 54 participants, whereas all fMRI analyses and brain–behaviour correlations are based on the 45 participants with usable imaging data.

All participants were right-handed and had normal hearing and normal or corrected-to-normal vision with no history of neurological or psychiatric disorders. They received monetary compensation for their participation. All participants gave written informed consent prior to participation. The study was approved by the Institutional Review Board of Korea University (approval no. 1040548-KU-IRB-13-42-A-2, approved on 31 March 2017), in accordance with the Declaration of Helsinki.

### 2.2. Stimuli and tasks

Participants performed two experimental tasks which involved making rhyming and spelling judgments on two words that appeared visually in a sequence. In rhyme judgement task, participants decided whether the two words rhymed in English. In spelling judgement task participants determined whether the two words had same spellings from the first vowel onwards (i.e., rime).

A list of 80 English word pairs was used for both tasks, with 20 pairs for each of four types that were manipulated in terms of orthographic (letters) and phonological (sounds) similarities between the words: (1) word pairs that have similar phonological but different orthographic rimes (P+O–, e.g., *light-kite*); (2) word pairs with different phonological but similar orthographic rimes (P–O+, e.g., *mint-pint*); (3) word pairs with similar phonological and orthographic rimes (P+O+, e.g., *bat-hat*); (4) word pairs with different phonological and orthographic rimes (P–O–, e.g., *glass-arm*). The first two types (P+O–, P–O+) constituted a conflict condition where orthographic and phonological information did not match, and the last two types (P+O+, P–O–) constituted a non-conflict condition where orthographic and phonological information matched so that the task difficulty level would be lower than the conflict condition. Throughout, conflict and non-conflict refer to the relationship between the orthographic and phonological rimes within the word pair, and this stimulus-level definition is identical for the two tasks. The conditions nevertheless map onto the two tasks in different ways. In the rhyme task, conflict pairs require the phonological decision to be made against competing orthographic evidence, whereas in the spelling task the same pairs require the orthographic decision to be made against competing phonological evidence. Conflict trials therefore combine stimulus-level orthographic–phonological conflict with response-level conflict, as they are also the trials on which the orthographically driven and the phonologically driven responses diverge; and for items with inconsistent grapheme-to-phoneme mappings (e.g., pint) the phonological form itself is in conflict, depending on whether it is assembled by grapheme-to-phoneme conversion or retrieved through lexical mediation. The words in all pairs were monosyllabic English words ranging from 3 to 6 letters, and were matched for log word frequency (*F*(3, 156) = 1.337, *p* = .264) and number of letters (*F*(3, 156) = 1.664, *p* = .177) across the types (Baayen et al., 1995) (Table 2).

**Table 2.**
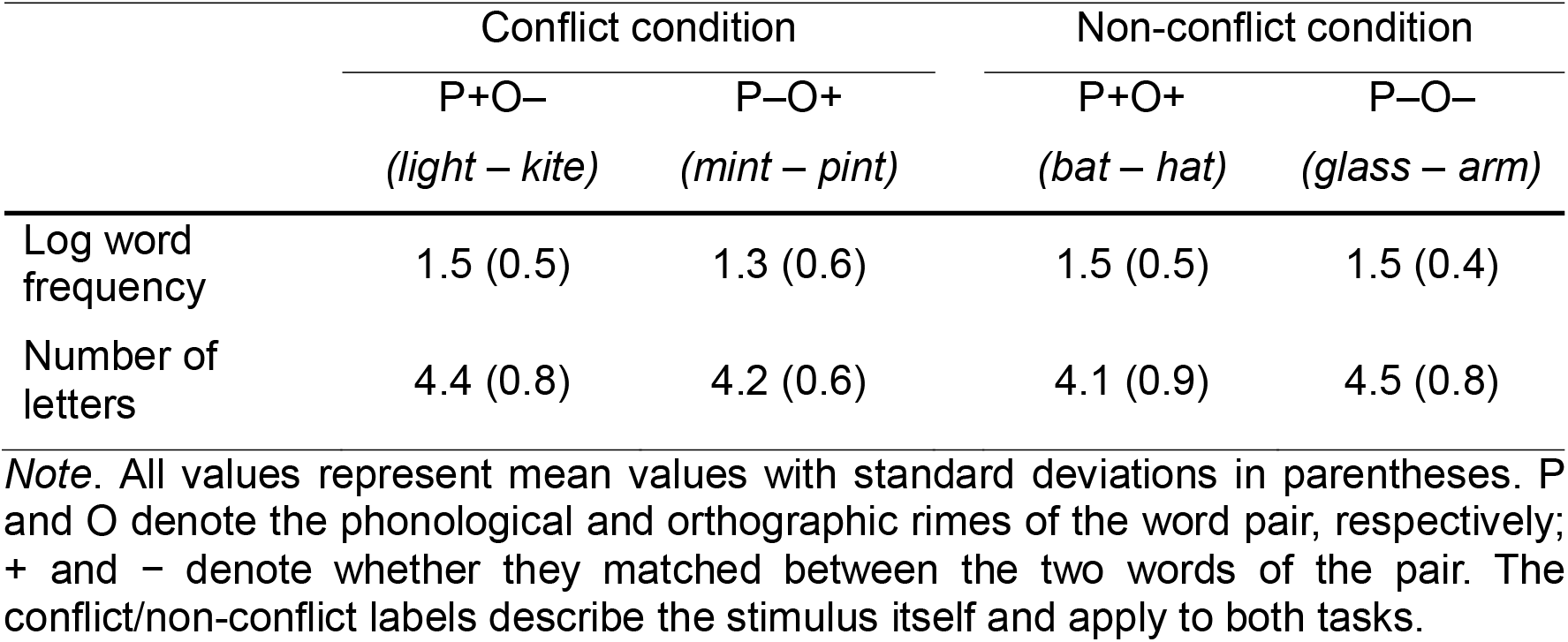
Stimuli characteristics for rhyme and spelling judgement tasks.

|  | Conflict condition |  | Non-conflict condition |  |
| --- | --- | --- | --- | --- |
|  | P+O–<br>(*light – kite*) | P–O+<br>(*mint – pint*) | P+O+<br>(*bat – hat*) | P–O–<br>(*glass – arm*) |
| Log word frequency | 1.5 (0.5) | 1.3 (0.6) | 1.5 (0.5) | 1.5 (0.4) |
| Number of letters | 4.4 (0.8) | 4.2 (0.6) | 4.1 (0.9) | 4.5 (0.8) |

*Note.* All values represent mean values with standard deviations in parentheses. P and O denote the phonological and orthographic rimes of the word pair, respectively; + and – denote whether they matched between the two words of the pair. The conflict/non-conflict labels describe the stimulus itself and apply to both tasks.

A control task was employed which required participants to press the “yes” button when a red fixation cross appeared on the screen, as opposed to a black fixation cross. Each fixation cross was presented for 800 ms, across a total of 80 trials. E-Prime 2.0 standard software (Psychology Software Tools, Pittsburgh, PA) was used for stimulus presentation and response collection. A complete stimuli list is provided in the **Supplementary Materials** (**Table S1**).

### 2.3. Procedure

Participants participated in the experimental scanning after the questionnaires and L2 proficiency measures. They engaged in a practice session to get familiarized with the rhyme and spelling judgement tasks before the actual experiment. Stimuli used during the practice session were not included in the experimental session. During scanning, the participants carried out the rhyme and the spelling judgement tasks in separate blocks. In both tasks, there were 80 trials lasting 6 seconds each. In a trial, each word of the pairs was displayed for 800 ms followed by a 200 ms blank screen (**Fig. 1**). A red fixation cross and a blank screen appeared after the second word. Participants then had to make a rhyming or spelling decision during the subsequent 3 second interval. They pressed a button with their left and right index fingers which corresponded to their judgement response. The order of stimuli within each task was fixed for all participants whereas the order of tasks was counterbalanced across participants.

**Figure 1.**
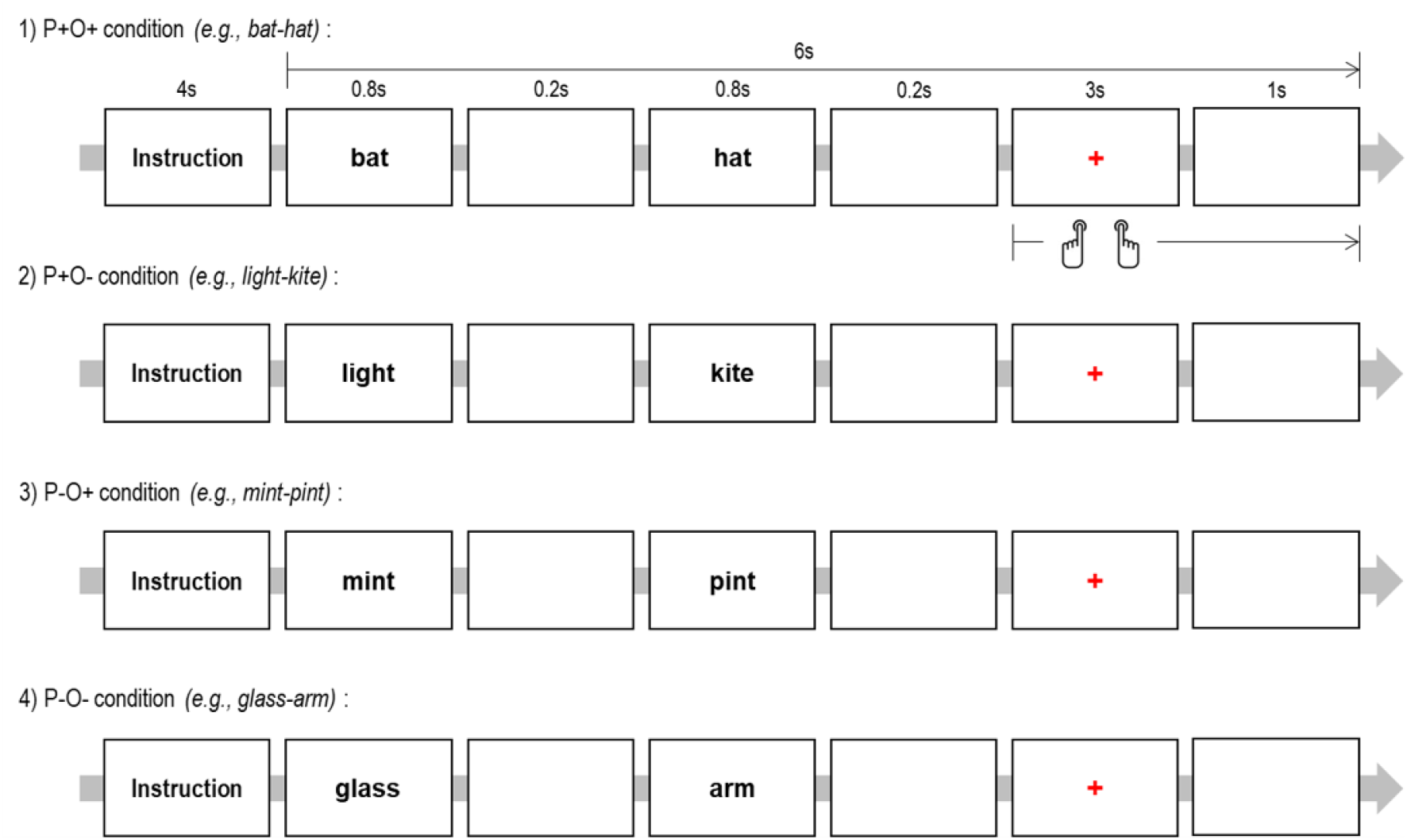
Experimental paradigm and trial structure. Schematic illustration of the rhyme and spelling judgement tasks performed in the scanner.

### 2.4. Image acquisition

All images were collected using a Siemens Magnetom Trio 3T MRI scanner at the Korea University Brain Imaging Centre, Seoul, South Korea. Structural images were acquired using Magnetization-Prepared Rapid Gradient-Echo (MPRAGE) sequence (TR = 1900 ms, TE = 2.52 ms, Flip Angle = 90°, Field of View = 256 mm, matrix size = 256 × 256, resolution = 1 mm × 1 mm × 1 mm). Functional images were acquired using a single-shot gradient Echo Planar Imaging (EPI) sequence with the following parameters: TR = 2000ms, TE = 20ms, Flip Angle = 90°, Field of View = 240 mm, slice thickness = 3 mm, 42 slices, matrix size = 80 × 80, and voxel size = 3 mm × 3 mm × 3 mm.

### 2.5. General linear model analysis

SPM12 (http://www.fil.ion.ucl.ac.uk/spm/software/spm12/) was used for image analysis. The first three functional volumes were discarded to reduce the transition effects of hemodynamic responses. The remaining images were realigned for motion correction, generating six rigid-body parameters. Participants who showed > 2 mm and/or 2° head motion were excluded from this study (*N* = 9). Then, the data were corrected for slice timing, co-registered with the structural images, then spatially normalised to a standard Montreal Neurological Institute (MNI) space and smoothed with an isotropic Gaussian kernel of 8 mm FWHM.

At the individual level, the data were modelled using general linear modelling (GLM). Conditions were modelled using a box-car function convolved with the canonical hemodynamic response function. Four separate regressors were modelled according to types (P+O–, P–O+, P+O+, P–O–) and head movement parameters estimated from the realignment were entered as additional regressors. At the group level, two-factorial ANOVA with condition (conflict: P+O–, P–O+, non-conflict: P+O+, P–O–) and group (advanced, intermediate, and beginner) was conducted for the main effect of task and group as well as interaction between condition and group for the rhyme judgement task and spelling judgement task. T-contrast were established for the contrast conflict > non-conflict, non-conflict > conflict according to the groups. Whole-brain maps were thresholded at *p* < .001 at the voxel level, with family-wise error (FWE)-corrected cluster threshold of *p* < .05, *k*s > 30.

### 2.6. Region of interest analysis

Region of interest (ROI) analysis was performed to investigate the group difference in the key region of visual word processing. Eight ROIs were selected based on the meta-analysis outcomes from relevant literatures related to reading (Martin et al., 2015) and phonological processing in visual words (Tan et al., 2005). The ROIs included the left inferior frontal gyrus (IFG) [MNI: –47 14 19], supplementary motor area (SMA) [MNI: –6 14 50], intraparietal sulcus (IPS) [MNI: –42 –48 48], inferior parietal lobe (IPL) [MNI: –55 –41 24], superior temporal gyrus (STG) [MNI: –56 –30 14], inferior temporal gyrus (ITG) [MNI: –48 –62 –20], fusiform gyrus (FG) [MNI: –44 –54 –12], and inferior occipital gyrus (IOG) [MNI: –44 –74 –4]. In addition, ROI analysis was conducted for the areas showing the interaction effect between the condition and group from the GLM analysis. All were defined as a spherical ROI with a radius of 6 mm.

### 2.7. Brain and behavioural relationship

In order to link bilinguals’ L2 proficiency and neural responses, we performed correlation analyses between the behavioural performance (task performance and TIWRE scores) and regional activity in the contrast of conflict > non-conflict (*p*_false discovery rate (FDR)-corrected_ < .05).

## 3. Results

### 3.1. Behavioural results

For each of the two tasks (rhyme judgement task, spelling judgement task), repeated measures ANOVAs were conducted separately on accuracy and RTs, with factors condition (conflict, non-conflict) and group (advanced, intermediate, beginner).

#### 3.1.1. Accuracy

For the rhyme judgement task, ANOVA showed a significant main effect of condition (*F*(1, 51) = 229.77, *p* < .001), with higher accuracy in the non-conflict (*M* = 95.1%, *SD* = 6.7) than in the conflict condition (*M* = 64.8%, *SD* = 16.4) (**Fig. 2A**). The main effect of group (*F*(2, 51) = 3.31, *p* = .04) and an interaction between group and condition were significant (*F*(2, 51) = 7.52, *p* = .001). Follow-up comparisons revealed that in the conflict condition, the advanced group performed significantly better than the beginner group (Bonferroni-corrected, *p* = .005; advanced: *M* = 73.4%, *SD* = 13.8; beginner: *M* = 56.0%, *SD* = 16.6), whereas all other group differences in both conditions were not significant (**Fig. 2A**). As noted earlier, the beginner group’s AoA was significantly older than that of the other groups. To control for this factor, we conducted an analysis of covariance (ANCOVA) with AoA as covariate. Results were overall consistent with those from the ANOVA. Specifically, the main effect of condition (*F*(1, 50) = 20.28, *p* < .001) and an interaction between group and condition remained significant (*F*(2, 50) = 5.61, *p* = .006), with follow-up comparisons showing higher accuracy for the advanced group than the beginner group in the conflict condition (Bonferroni-corrected, *p* = .026).

**Figure 2.**
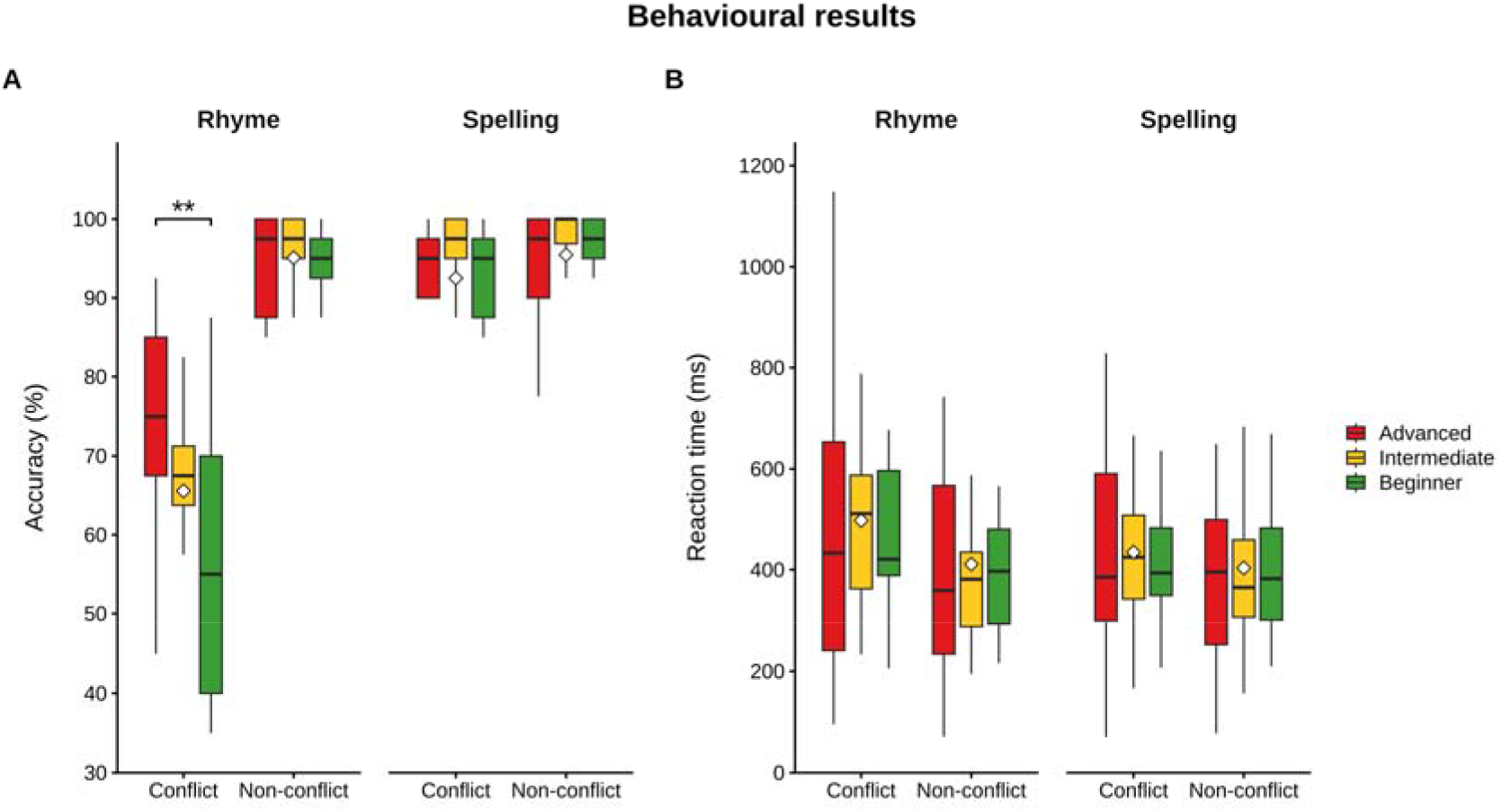
Behavioural performance in the rhyme and spelling judgement tasks. (A) Accuracy (%) and (B) reaction time (ms) for the advanced, intermediate, and beginner groups in the conflict and non-conflict conditions. Within each panel, the rhyme judgement task is shown on the left and the spelling judgement task on the right. Boxes indicate the median and interquartile range, whiskers extend to 1.5× the interquartile range, and white diamonds mark group means. Asterisks denote significant pairwise post-hoc comparisons (Bonferroni-corrected, *p* < .01).

For the spelling judgement task, ANOVA showed a significant main effect of condition (*F*(1, 51) = 6.39, *p* = .015), with higher accuracy in the non-conflict (*M* = 95.5%, *SD* = 7.2) than in the conflict condition (*M* = 92.5%, *SD* = 11.3) (**Fig. 2A**). Neither the main effect of group (*F*(2, 51) = 2.71, *p* = .076) nor the interaction between group and condition (*F*(2, 51) = 0.76, *p* = .475) was significant. ANCOVA with AoA as a covariate showed no significant effects.

Correlations were computed between TIWRE scores and accuracy for each task and condition, yielding four correlations. Bonferroni correction was applied to correct for multiple comparisons. TIWRE scores were significantly positively correlated with accuracy in the rhyme judgement task under the conflict condition (r = 0.62, corrected *p* < 0.004), whereas no association was found in the non-conflict condition (*r* = 0.00, corrected *p* = 1.00). Partial correlations controlling for AoA showed similar results, revealing a significant positive correlation between TIWRE scores and accuracy in the same condition (r = 0.59, corrected *p* < 0.004), with again no association in the non-conflict condition (*r* = –0.003, corrected *p* = 1.00).

As the two conflict types are not equivalent, we also inspected the sub-conditions of the rhyme task. Accuracy was markedly lower for P–O+ pairs (e.g., mint–pint; *M* = 51.4%, *SD* = 23.6) than for P+O– pairs (e.g., light–kite; *M* = 78.2%, *SD* = 16.1; *t*(53) = 8.34, *p* < .001), and the P–O+ sub-condition showed the clearest proficiency grading (advanced: *M* = 63.2%, *SD* = 24.4; intermediate: *M* = 52.2%, *SD* = 20.0; beginner: *M* = 38.5%, *SD* = 21.3) as well as the strongest association with L2 reading ability (*r* = 0.68 with TIWRE, versus *r* = 0.26 for P+O– and *r* < 0.03 for the two non-conflict sub-conditions). The conflict effect in the rhyme task is therefore carried largely by pairs whose shared spelling masks a difference in pronunciation.

#### 3.1.2. Reaction Times

For the rhyme judgement task, there was only a significant main effect of condition (*F*(1, 51) = 17.73, *p* < .001), with slower RTs in the conflict than non-conflict conditions (conflict: *M* = 497 ms, *SD* = 237; non-conflict: *M* = 411 ms, *SD* = 179; **Fig. 2B**). ANCOVA controlling for AoA revealed no significant effects.

For the spelling judgement task, there was a significant main effect of condition (*F*(1, 51) = 14.51, *p* < .001), with slower RTs in the conflict than non-conflict condition (conflict: *M* = 435 ms, *SD* = 181; non-conflict: *M* = 403 ms, *SD* = 169; **Fig. 2B**). The interaction between group and condition was also significant (*F*(2, 51) = 3.32, *p* = .044), but follow-up comparisons showed no significant group differences in either the conflict (*F*(2, 51) = 0.17, *p* = .845) or the non-conflict condition (*F*(2, 51) = 0.72, *p* = .491). No significant effects were observed in ANCOVA.

Correlations were computed between TIWRE scores and RTs for each task and condition, with Bonferroni correction applied. No significant correlations were observed (all corrected *p*s > .10), and partial correlations controlling for AoA likewise revealed no significant correlations (all corrected *p*s > .31).

### 3.2. fMRI results

In the rhyme judgement task, the whole brain analysis revealed that the conflict condition evoked significant activation in the bilateral IFG, SMA, middle cingulate cortex (MCC), left pre/postcentral gyrus, left insula, bilateral supramarginal gyrus (SMG), left IPL, left superior parietal lobe (SPL), right angular gyrus (AG), caudate nucleus, left ITG, right cerebellum, and visual cortex, whereas the non-conflict condition revealed activation in the superior frontal gyrus (SFG), middle orbital gyrus (MOG), STG, right pre/postcentral gyrus, and left AG across the groups (**Fig. 3A**). The results demonstrated that bilinguals recruited the central executive control system - frontoparietal network (FPN) for demanding L2 processing (conflict) by suppressing the default mode network (DMN). Specifically, the advanced group showed increased activation in the FPN as well as the left posterior MTG during the conflict condition relative to the non-conflict condition. The intermediate group exhibited more widespread activation in the FPN and deactivation in the superior medial gyrus in the contrast of conflict > non-conflict. The beginner group showed the left-lateralized activation in the frontoparietal regions without caudate activation during the conflict condition. The advanced and beginner groups did not show any activation in the contrast of non-conflict > conflict. The results are summarised in **Table S2**.

**Figure 3.**
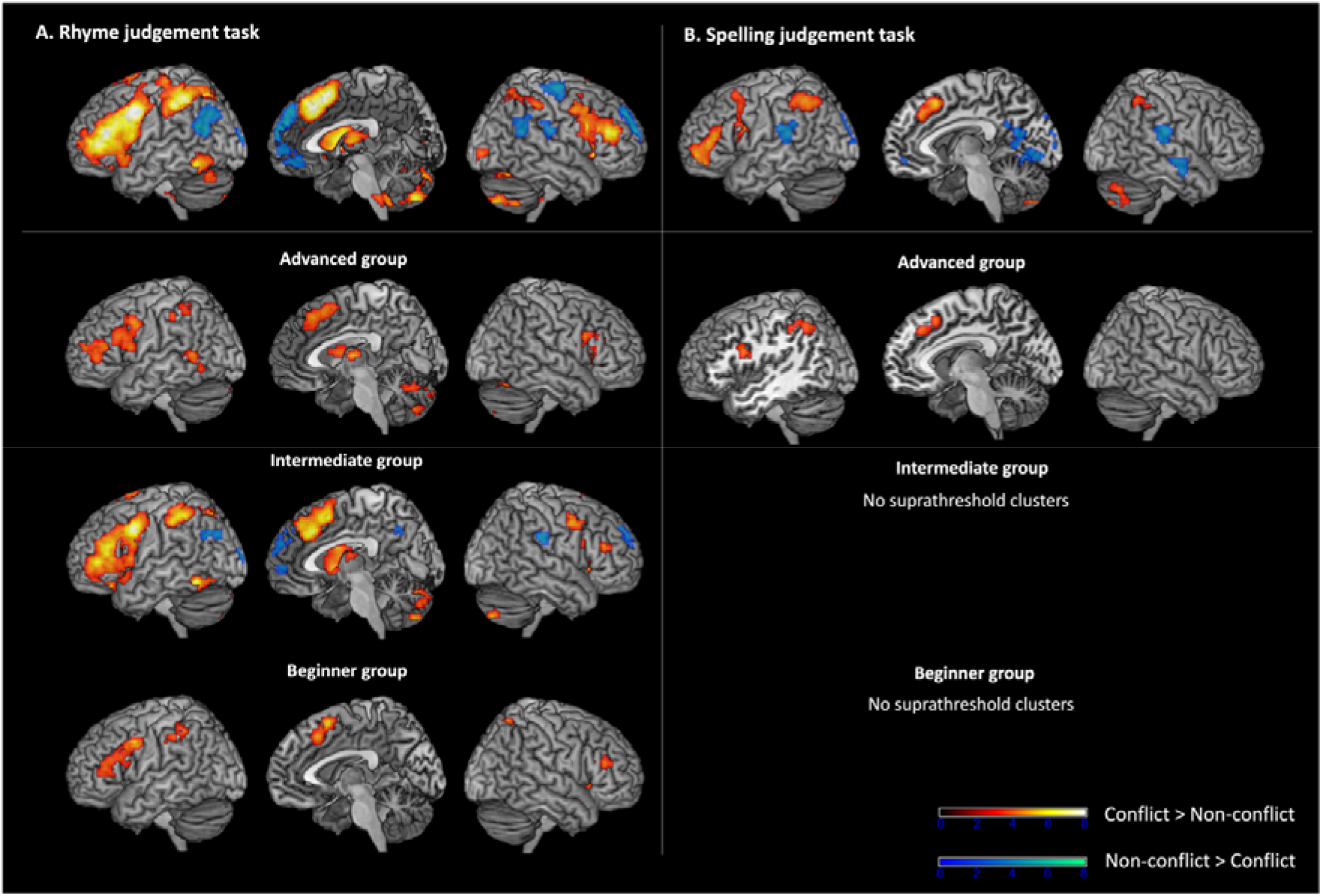
Whole-brain activation for conflict versus non-conflict trials in each task. (A) Rhyme judgement task. (B) Spelling judgement task. In each panel, the top row shows the whole imaging sample (*N* = 45) and the lower rows the advanced (*N* = 16), intermediate (*N* = 14), and beginner (*N* = 15) groups. Maps are thresholded at a voxel-level *p* < 0.001 with a cluster-level FWE-corrected *p* < 0.05. Warm colours indicate conflict > non-conflict, cool colours indicate non-conflict > conflict. Colour bars show t-values. No suprathreshold clusters were observed for the intermediate or beginner groups in the spelling judgement task.

In the spelling judgement task, the conflict condition compared to non-conflict condition induced activation in the left IFG, SMA, bilateral IPL, and right cerebellum as well as deactivation in the bilateral lingual gyrus, calcarine gyrus, precuneus, posterior cingulate cortex (PCC), cuneus, STG, and right SMG (**Fig. 3B**). Only advanced group showed the increased activation in the left IFG and IPL in the contrast of conflict > non-conflict. There was no significant difference between the conditions in the intermediate and beginner groups. The results are summarised in **Table S3**.

Critically, several regions showed a condition by group interaction, specific to each task (**Fig. 4**). The results are summarised in **Table S4**. In the rhyme judgement task, significant interactions were observed in the left SMG, middle orbital gyrus (MOG), dorsal medial prefrontal cortex (dmPFC), and cerebellum (**Fig. 4A**). To characterise the direction of these interactions, we extracted the conflict > non-conflict contrast estimate from each ROI and compared groups using one-way ANCOVA with AoA as a covariate. Contrast estimates differed across groups in all four ROIs (SMG: *F*(2, 41) = 3.95, *p* = .027, MOG: *F*(2, 41) = 4.14, *p* = .023, dmPFC: *F*(2, 41) = 4.68, *p* = .015, cerebellum: *F*(2, 41) = 6.32, *p* = .004). Post-hoc t-tests with Bonferroni correction for multiple comparisons revealed that in the left SMG and the cerebellum, estimates were highest in the advanced group. For the SMG, this held relative to both the intermediate (*p* = .036) and the beginner group (*p* = .043). For the cerebellum, the difference from the beginner group was significant (*p* = .004), whereas the difference from the intermediate group did not reach for the corrected threshold (*p* = .07). The left MOG showed the opposite ordering, with the highest estimates in the intermediate group, which was significant relative to the beginner group (*p* = .045), but not relative to the advanced group (*p* = .063). The intermediate group also showed higher dmPFC estimates than the beginner group (*p* = .016).

**Figure 4.**
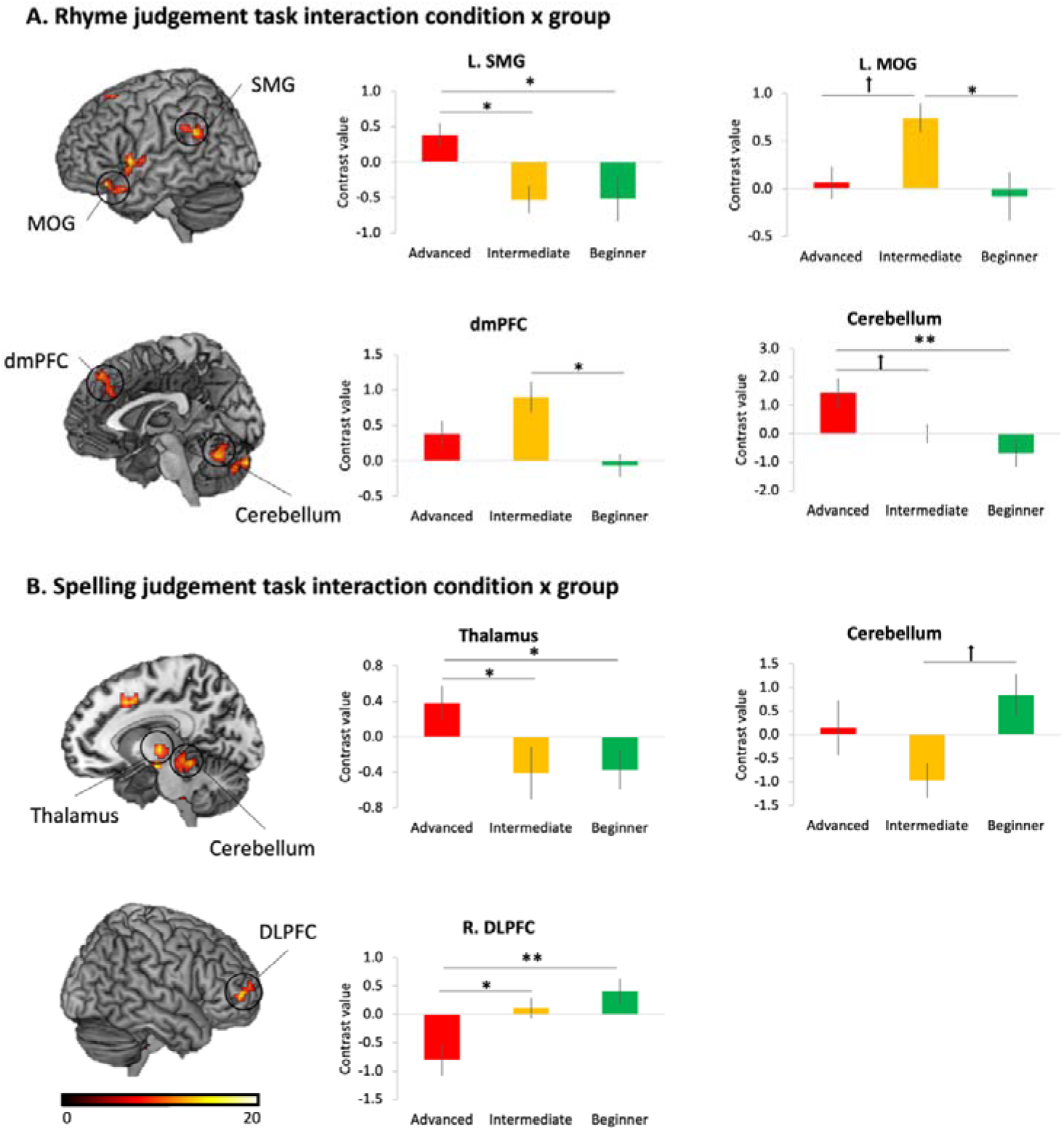
Regions showing a group × condition interaction. ROIs in which the conflict > non-conflict contrast varied significantly by proficiency group, shown separately for each task. Bar plots present mean contrast estimates (± SEM) for the advanced, intermediate, and beginner groups, with AoA included as a covariate. (A) Rhyme judgement task: left supramarginal gyrus (SMG), left middle orbital gyrus (MOG), dorsomedial prefrontal cortex (dmPFC), and right cerebellum. (B) Spelling judgement task: thalamus, left cerebellum, and right dorsolateral prefrontal cortex (DLPFC). \*\**p* < .01; \**p* < .05; ^†^*p* < .08;

In the spelling judgement task, three regions showed the interaction effect including the thalamus, left cerebellum, and right dorsolateral prefrontal cortex (DLPFC) (**Fig. 4B**). Contrast estimates differed across groups in the thalamus (*F*(2, 41) = 4.63, *p* = .015) and the right DLPFC (*F*(2, 41) = 6.32, *p* = .004). There was a marginally significant group effect in the cerebellum (*F*(2, 41) = 2.81, *p* = .072). Post-hoc t-tests (Bonferroni corrected) demonstrated that the advanced group had strengthened regional activity in the thalamus compared to the intermediate (*p* = .044) and beginner (*p* = .032) groups. The cerebellum showed the stronger activation in the beginner group than the intermediate group (*p* = .075). The right DLPFC revealed deactivation in the advanced group compared to the intermediate (*p* = .030) and beginner (*p* = .006) groups.

We subsequently focused on the eight a priori ROIs associated with visual word processing to examine the effect of L2 proficiency responding to task conditions (**Fig. 5**). ANOVA with task (rhyme vs. spelling) and condition (conflict vs. non-conflict) as a within subject factor and group (advanced, intermediate, beginner) as a between subject factor was conducted for each ROI. The SMA revealed the significant main effect of condition (*F*(1, 42) = 7.60, *p* = .009). The IFG showed the main effect of condition (*F*(1, 42) = 13.99, *p* < .001). The IPS revealed the main effect of condition (*F*(1, 42) = 6.61, *p* = .014). The FG revealed the main effect of task (*F*(1, 42) = 5.71, *p* = .022). The IPL, STG, ITG, and IOG did not show any effects. The results demonstrated that the key regions of visual word processing did not show L2 proficiency effects in L2 processing.

**Figure 5.**
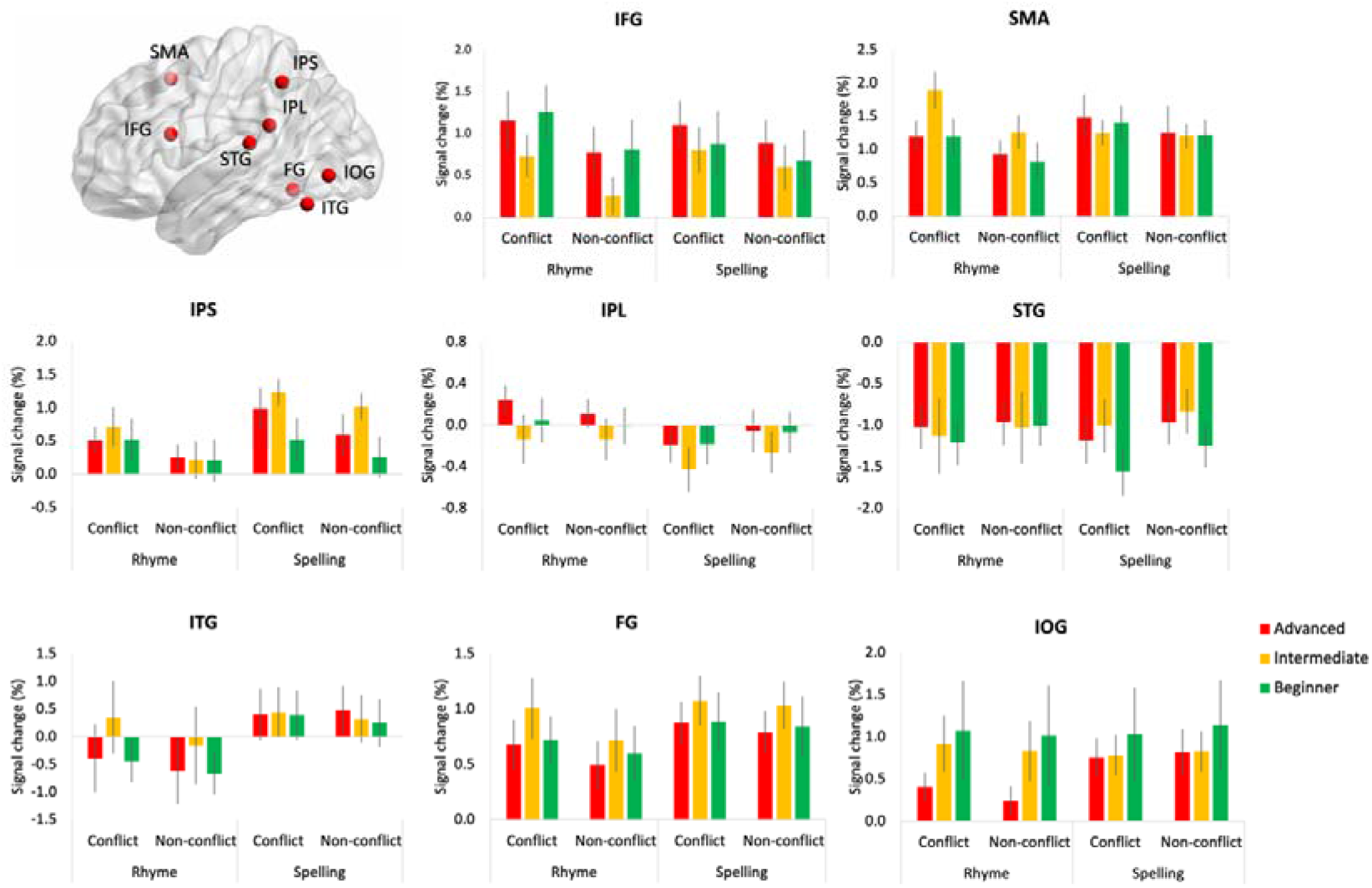
ROI activation in canonical visual word recognition regions. Mean BOLD response (± SEM) extracted from a priori ROIs implicated in visual word processing, supplementary motor area (SMA), inferior frontal gyrus (IFG), intraparietal sulcus (IPS), fusiform gyrus (FG), inferior parietal lobule (IPL), superior temporal gyrus (STG), inferior temporal gyrus (ITG), and inferior occipital gyrus (IOG), plotted as a function of task (rhyme, spelling), condition (conflict, non-conflict), and group (advanced, intermediate, beginner).

### 3.3. Relationship between L2 performance and fMRI neural activation

In order to investigate how L2 proficiency influences neural response in bilinguals, correlation analyses were conducted between bilinguals’ L2 performance (task performance: accuracy and TIWRE score) and the regional BOLD response (the conflict > non-conflict contrast estimate extracted from each ROI) in the areas showing the interaction effect between the condition and group (**Fig. 4**). Rhyme accuracy in the conflict condition was significantly correlated with regional activity in the cerebellum (*r* = 0.37, *p*_FDR-corrected_ = .031) and showed a non-significant trend in the left SMG (*r* = 0.30, *p*_FDR-corrected_ = .10) in the rhyme judgement task (**Fig. 6A**). The regional activity in these regions was positively correlated with TIWRE score (cerebellum: *r* = 0.41, *p*_FDR-corrected_ = .012; SMG: *r* = 0.36, *p*_FDR-corrected_ = .049) (**Fig. 6B**). Bilinguals with stronger activity in the cerebellum and left SMG performed better in the rhyme judgement task and had higher scores in TIWRE. In contrast, bilinguals with stronger activity in the right DLPFC showed the lower TIWRE score (*r* = –0.35, *p*_FDR-corrected_ = .031).

**Figure 6.**
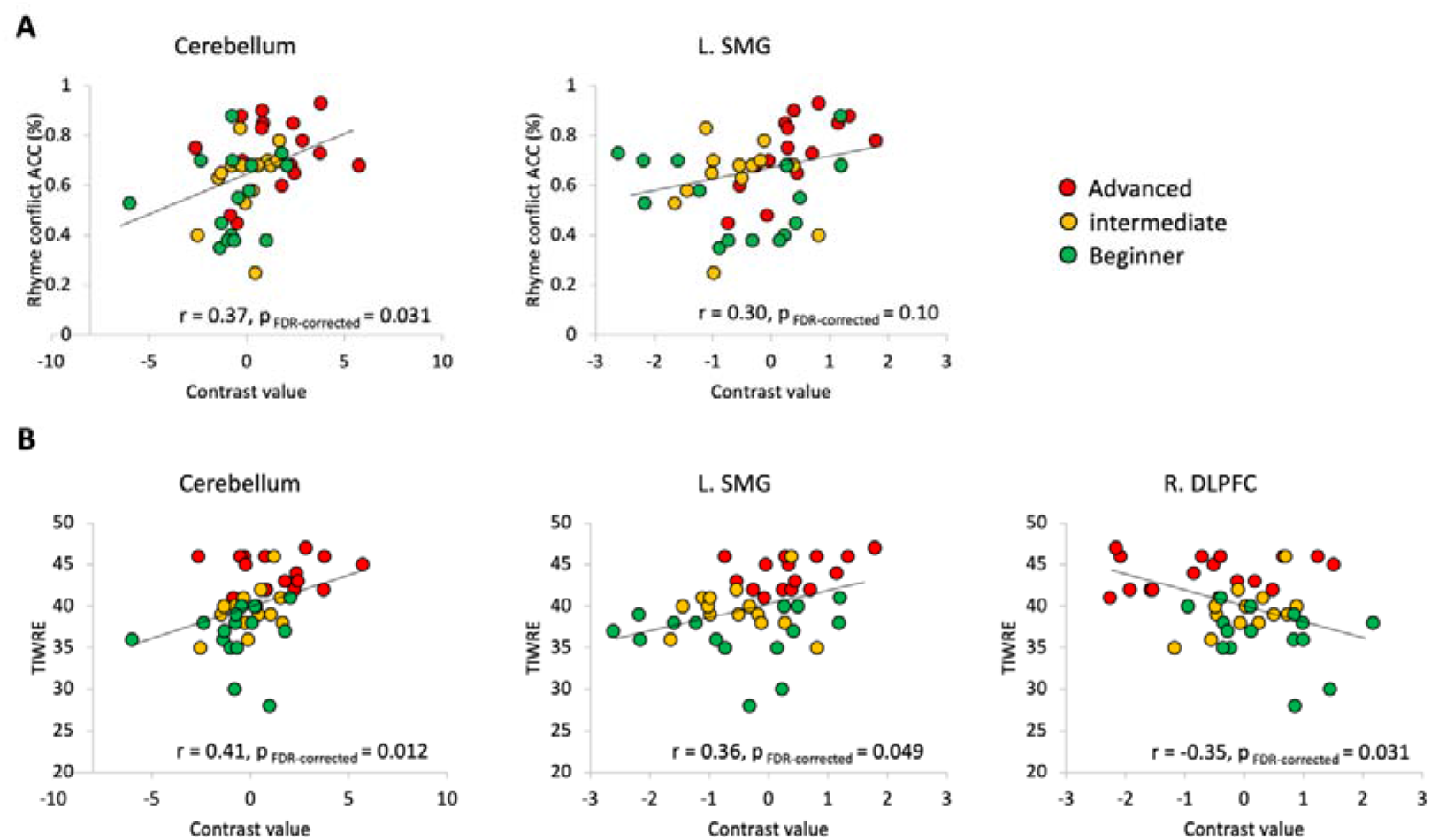
Relationship between the conflict > non-conflict contrast estimate and behavioural measures of L2 reading. Scatterplots relating regional activity in the interaction-effect ROIs (Fig. 4) to behavioural indices of L2 reading ability. (A) The conflict > non-conflict contrast estimate (x-axis) for the right cerebellum and left SMG in the rhyme judgement task plotted against accuracy in the conflict condition of that task (y-axis). (B) The same contrast estimates for those regions, together with the right DLPFC, plotted against standardised L2 reading scores (TIWRE). Each point represents an individual participant; regression lines with 95% confidence intervals are overlaid.

## 4. Discussion

The present study examined how L2 proficiency modulates the neural activity during L2 English reading in Korean–English (K–E) bilinguals across rhyme and spelling judgement tasks involving orthographic-phonological conflict. Canonical reading regions showed relatively limited proficiency-dependent modulation, with no significant group differences in conflict-related activation. Instead, proficiency effects were observed in regions associated with cognitive control, attentional regulation, phonological storage, and motor-articulatory prediction, including the cerebellum, SMG, thalamus, and lateral and medial prefrontal cortex. During the rhyme judgement task, advanced bilinguals exhibited greater activation in the left supramarginal gyrus (SMG) and right cerebellum, regions implicated in grapheme-to-phoneme conversion, phonological storage, and articulatory rehearsal (Booth et al., 2007; Stoeckel et al., 2009; Stoodley and Schmahmann, 2009). Importantly, activity in both regions was positively associated with task performance and standardised L2 reading ability (TIWRE), suggesting that higher proficiency may be supported by more efficient recruitment of phonological and predictive-articulatory systems during conflict resolution in the L2. In contrast, intermediate bilinguals showed greater activation in the left middle orbital gyrus (MOG) extending into anterior inferior frontal cortex, as well as dorsomedial prefrontal cortex (dmPFC), regions commonly associated with conflict monitoring and effortful cognitive control (Botvinick et al., 2004; Niendam et al., 2012). A different proficiency-related pattern emerged with the spelling judgement task. Advanced bilinguals demonstrated greater thalamic activation, consistent with enhanced attentional gating and coordination of distributed reading systems, alongside greater deactivation of the right dorsolateral prefrontal cortex (DLPFC), suggesting reduced reliance on domain-general executive control during orthographic processing.

Together, these results suggest that variation in L2 reading proficiency among K-E bilinguals may be characterised less by changes within the core language network *per se* than by task-specific shifts in the balance between phonological-specialised systems, subcortical attentional support, and domain-general control mechanisms during L2 processing.

### 4.1. Rhyme judgement under orthographic–phonological conflict recruits proficiency-dependent cerebellar and inferior parietal systems

The rhyme judgement task placed substantial demand on phonological processing, particularly under conflict situations where orthographic and phonological information were incongruent. As described in the Methods, conflict in this task operates simultaneously at the stimulus and response levels and, for items with inconsistent grapheme-to-phoneme mappings, at the level of the phonological code itself. The observed activation should therefore not be attributed to any single level. Indeed, conflict trials elicited robust activation across the FPN/ECN, including bilateral inferior frontal gyrus (IFG), supplementary motor area (SMA), mid-cingulate cortex, insula, inferior parietal lobe (IPL), and SMG, alongside DMN deactivation. These results are in line with previous studies of alphabetic reading showing that phonological mismatches recruit a distributed control system to resolve competition between orthographic and phonological information (Booth et al., 2004; Tan et al., 2005).

Activation in the right cerebellum and the left SMG showed a clear proficiency-dependent modulation. Advanced bilinguals exhibited stronger activation in both regions during conflict, and the activity in these regions was significantly correlated with the TIWRE score. The cerebellum has been consistently implicated in articulatory rehearsal, internal phonological prediction, and the timing of phonological computations during reading (Booth et al., 2007; Stoodley and Schmahmann, 2009), whereas the left SMG is a core hub for grapheme-to-phoneme conversion, phonological short-term storage, and rhyme-based decisions (Stoeckel et al., 2009; Tan et al., 2005). Cerebellar activity was also positively associated with rhyme accuracy in conflict trials, whereas the parallel correlation between left SMG activity and rhyme accuracy was only a trend and should therefore be interpreted with caution. Taken together, these patterns are consistent with the view that proficient K-E bilinguals may resolve orthographic-phonological conflict in the L2 by placing greater weight on phonologically specialized and articulatory-predictive resources, alongside the shared recruitment of frontoparietal control systems.

Intermediate bilinguals exhibited a distinct pattern, characterised by selectively elevated activation in the left mid-orbital gyrus (extending into the anterior IFG) and the dorsomedial prefrontal cortex (dmPFC). Unlike classical language regions, these prefrontal areas have been linked to effortful conflict monitoring, performance evaluation, and self-referential processing (Andrews-Hanna, 2012; Botvinick et al., 2004). One explanation is that intermediate proficiency reflects a transitional stage in which orthographic-phonological conflict places maximal demands on top-down monitoring systems. In contrast, advanced bilinguals may process conflict more efficiently through phonological-specialized systems, whereas beginners may engage these monitoring systems less consistently. This non-linear pattern broadly resembles previous reports suggesting heightened control-system recruitment during intermediate stages of L2 development (Wang et al., 2020).

### 4.2. Orthographic processing reveals proficiency-related differences despite similar behavioural performance

Although the spelling judgement task showed limited behavioural differences across proficiency groups, neural activation differed systematically as a function of proficiency. This dissociation suggests that orthographic decisions in L2 English continue to engage proficiency-sensitive neural mechanisms in K-E bilinguals, even when behaviour is comparable across groups, in line with previous reports that bilingual neural differences can emerge in the absence of behavioural differences (Cargnelutti et al., 2019; Wang et al., 2020).

Advanced bilinguals exhibited stronger thalamic activation on conflict trials than the intermediate and beginner groups. The thalamus is a key node for attentional gating and cortico-cortical communication and has also been implicated in reading and bilingual language control (Burgaleta et al., 2016; Cao et al., 2013). The present pattern is consistent with the proposal that L2 expertise may be associated with greater subcortical support for cortical reading networks, which may have contributed to the relatively stable behavioural performance of advanced bilinguals on spelling decisions.

In contrast, beginners showed relatively greater cerebellum activation during the spelling task. As the Korean writing system has highly transparent letter-to-sound mappings, beginners may rely more strongly on subvocal phonological strategy when processing English orthography, thereby increasing demands on articulatory rehearsal systems involving cerebellum (Booth et al., 2007; Stoodley and Schmahmann, 2009). Notably, the cerebellum exhibited opposite proficiency patterns across tasks, with greater activation in advanced bilinguals during phonological conflict but greater activation in beginners during orthographic processing. This dissociation suggests that cerebellar contributions to L2 reading are both task-and proficiency-dependent.

The right dorsolateral prefrontal cortex (DLPFC) showed graded modulation across proficiency levels, with advanced group exhibiting relative deactivation and intermediate and beginner groups showing progressively higher activation. Right DLPFC has been widely implicated in domain-general conflict monitoring, response inhibition, and effortful control (Niendam et al., 2012), and bilingual neuroimaging work has linked greater right-prefrontal engagement to less proficient or less automatic L2 use (Abutalebi and Green, 2007). Crucially, right DLPFC activity was negatively correlated with the TIWRE score, suggesting that increasing proficiency may reduce reliance on effortful domain-general control system during orthographic processing.

### 4.3. Proficiency-dependent engagement of language and control system

Taken together, the present findings support the view that increasing L2 proficiency alters the relative engagement of language-related and control systems during L2 processing. Lower proficiency was associated with stronger reliance on right prefrontal control system and cerebellar resources during orthographic decisions, consistent with a more effortful and phonologically mediated approach to L2 English word processing. Intermediate proficiency was characterised by increased recruitment of prefrontal monitoring regions (mid-orbital gyrus, dmPFC), whereas advanced proficiency was associated with greater engagement of phonologically specialized cortical hubs (left SMG), motor-articulatory predictive systems (cerebellum), and subcortical attentional regulators (thalamus).

These findings broadly align with neurobiological models proposing that L2 development involves shifting interactions between domain-general and language-specific systems. The Adaptive Control Hypothesis (ACH; Green and Abutalebi, 2013) predicts that bilingual language processing dynamically recruits cognitive control system according to task demands, whereas the Dynamic Restructuring Model (DRM; DeLuca et al., 2019; Pliatsikas, 2020) and the Bilingual Anterior to Posterior and Subcortical Shift (BAPSS; Grundy et al., 2017) framework propose increasing involvement of posterior and subcortical systems with greater L2 experience and automaticity. The present findings are consistent with these accounts, although the absence of strong group differences in canonical left-lateralised reading network suggests that proficiency effects in the current dataset are better characterised as differential engagement of supporting control, attention, and prediction systems, rather than as a wholesale reorganisation of core language representations.

The present findings can also reflect the distinct orthographic relationship between Korean and English. Korean supports relatively transparent grapheme– phoneme mapping, whereas English requires more flexible and more lexically mediated decoding strategies. K-E bilinguals may therefore need to progressively downregulate L1-driven phonological decoding strategies while developing more efficient coordination between phonological, predictive, and attentional systems during L2 English reading. This interpretation is broadly consistent with prior work on K-E bilinguals showing that L2 effects often manifest as changes in network coupling and supporting-system recruitment rather than as activation differences within canonical language regions (Jung et al., 2018).

### 4.4. Methodological considerations and limitations

Several limitations should be considered when interpreting the present findings. First, our sample size is relatively small and predominantly female (43 of 54 participants). Although sex composition did not differ across the three groups (Fisher’s exact p = .836), the study was not designed or powered to examine sex or gender as a factor, and we did not analyse it. It thus remains untested whether the proficiency-related patterns reported here generalise to a more balanced sample. Second, the cross-sectional design precludes strong conclusions about the developmental trajectory of L2 acquisition. The proficiency-related differences observed here should therefore be interpreted as cross-sectional contrasts that are consistent with, but do not directly demonstrate, within-individual reorganisation. Longitudinal designs in which the same learners are scanned across proficiency stages would more directly test the proposed trajectory. Third, the present findings are situated in the specific typological context of K-E bilingualism, in which an L1 with a shallow, alpha-syllabic script is paired with a deep alphabetic L2. Replications in bilinguals with other L1s, including alphabetic L1s that differ from English in orthographic depth, such as French or Spanish, as well as logographic L1s, would help clarify which features of the present pattern generalise. Studies of language pairs with smaller differences in orthographic depth would also be informative. Finally, the present design did not include an L1 English control group. We therefore cannot establish which components of the conflict effect are specific to L2 reading and which would also be observed in native English readers performing the same tasks. A monolingual comparison group would be needed to distinguish these possibilities.

## 5. Conclusion

The present study provides evidence that variation of L2 proficiency in Korean-English bilinguals is associated with graded differences in the engagement of neural systems supporting L2 English reading. Higher proficiency was associated with enhanced recruitment of phonologically specialized and predictive-articulatory systems, including the left SMG, cerebellum, thalamus, and reduced engagement of right-prefrontal control regions. In contrast, the core reading regions remained comparably stable across proficiency levels. These findings support the view that L2 proficiency may modulate the interaction between language-related and domain-general systems rather than simply altering activation within the core reading network itself.

## CRediT authorship contribution statement

**Joonwoo Kim**: Data curation, Formal analysis, Visualization, Writing – original draft. **Jiyoun Choi**: Methodology, Formal analysis, Writing – original draft. **Yeonji Baik**: Conceptualization, Investigation, Writing – original draft. **Walter J. B. van Heuven**: Conceptualization, Writing – review and editing. **Kichun Nam**: Conceptualization, Resources, Supervision. **JeYoung Jung**: Methodology, Formal analysis, Validation, Visualization, Supervision, Project administration, Writing – review and editing.

## Data availability

The behavioural data, region-of-interest contrast estimates, and analysis code supporting the findings of this study will be deposited on the Open Science Framework (OSF) and made openly available upon publication. In the interim, the data and code are available from the corresponding author on reasonable request.

