## Supplementary material for "Proficiency-Dependent Reorganisation of Language and Control Networks during Second Language Processing: An fMRI Study of Korean–English Bilinguals": Table S1 (from another version of this preprint: version 1, 2026-08-19)

**Supplementary table 1**

|  | Non-conflict condition | | | | | | Conflict condition | | | | | |
| --- | --- | --- | --- | --- | --- | --- | --- | --- | --- | --- | --- | --- |
|  | P+O+ | | | P-O- | | | P+O- | | | P-O+ | | |
| 1 | bag | – | flag | bride | – | ghost | bear | – | hair | bomb | – | comb |
| 2 | bat | – | hat | brush | – | skirt | bread | – | bed | bough | – | tough |
| 3 | beach | – | leach | bus | – | knight | cane | – | chain | bowl | – | owl |
| 4 | bell | – | shell | couch | – | tent | coat | – | note | break | – | leak |
| 5 | cake | – | snake | cup | – | fence | drill | – | heel | cow | – | row |
| 6 | clown | – | crown | desk | – | truck | ear | – | deer | dead | – | lead |
| 7 | fan | – | pan | fish | – | tank | fly | – | eye | done | – | tone |
| 8 | fox | – | box | glass | – | arm | key | – | tree | dough | – | cough |
| 9 | fridge | – | bridge | leg | – | cage | light | – | kite | dread | – | bead |
| 10 | frog | – | dog | lips | – | pool | melt | – | dealt | gear | – | wear |
| 11 | goat | – | boat | pipe | – | scarf | peach | – | switch | mint | – | pint |
| 12 | gun | – | sun | rain | – | girl | scale | – | snail | pear | – | near |
| 13 | house | – | mouse | rock | – | nurse | shoe | – | screw | pour | – | tour |
| 14 | nose | – | rose | shark | – | belt | sled | – | thread | span | – | swan |
| 15 | phone | – | bone | sink | – | slide | square | – | chair | steak | – | peak |
| 16 | skirt | – | shirt | smoke | – | book | tail | – | whale | stove | – | glove |
| 17 | spoon | – | moon | stairs | – | plate | thumb | – | drum | tread | – | mead |
| 18 | star | – | car | sword | – | bricks | train | – | plane | vow | – | low |
| 19 | vest | – | nest | teeth | – | tie | type | – | wipe | watch | – | match |
| 20 | wing | – | ring | witch | – | duck | wheel | – | seal | zone | – | gone |

**Table S1.** Stimuli List for Rhyme Judgement Task and Spelling Judgement Task
