## Supplementary material for "Proficiency-Dependent Reorganisation of Language and Control Networks during Second Language Processing: An fMRI Study of Korean–English Bilinguals": Table S2 (from another version of this preprint: version 1, 2026-08-19)

**Supplementary table 2**

| contrast | Region | cluster | MNI coordinate | | | T |
| --- | --- | --- | --- | --- | --- | --- |
|  |  |  | x | y | z |  |
| All | | | | | | |
| conflict > non-conflict | L Supplementary Motor Area | 4756 | -3 | 11 | 56 | 10.76 |
|  | L Precentral Gyrus |  | -51 | 5 | 35 | 9.98 |
|  | L IFG (p. Triangularis) |  | -42 | 35 | 17 | 8.83 |
|  | R MCC |  | 6 | 29 | 38 | 8.32 |
|  | L IFG (p. Triangularis) |  | -45 | 20 | 23 | 8.12 |
|  | L Caudate Nucleus |  | -12 | 11 | 5 | 8.08 |
|  | L Insula Lobe |  | -30 | 26 | -1 | 7.33 |
|  | L Inferior Parietal Lobule | 1150 | -45 | -43 | 50 | 7.84 |
|  | L Inferior Parietal Lobule |  | -36 | -43 | 41 | 7.63 |
|  | L Inferior Parietal Lobule |  | -48 | -34 | 44 | 7.63 |
|  | L Superior Parietal Lobule |  | -24 | -61 | 44 | 7.33 |
|  | L Precentral Gyrus |  | -30 | -25 | 68 | 4.77 |
|  | L Postcentral Gyrus |  | -45 | -25 | 53 | 4.32 |
|  | R Cerebellum (VIII) | 630 | 21 | -73 | -49 | 6.78 |
|  | R Cerebellum (VI) |  | 24 | -67 | -25 | 6.42 |
|  | R Cerebellum (Crus 2) |  | 33 | -76 | -49 | 5.4 |
|  | R Cerebellum (Crus 1) |  | 12 | -85 | -28 | 5.29 |
|  | R Angular Gyrus | 398 | 30 | -64 | 47 | 6.17 |
|  | R SupraMarginal Gyrus |  | 48 | -34 | 44 | 4.63 |
|  | R Inferior Parietal Lobule |  | 39 | -49 | 44 | 4.6 |
|  | R Middle Occipital Gyrus |  | 30 | -67 | 32 | 4.11 |
|  | L Inferior Temporal Gyrus | 176 | -54 | -52 | -16 | 6.68 |
|  | R Calcarine Gyrus | 116 | 15 | -73 | 8 | 4.61 |
| non-conflict > conflict | R Superior Frontal Gyrus | 310 | 15 | 59 | 23 | 6.47 |
|  | R Mid Orbital Gyrus |  | 3 | 44 | -10 | 4.2 |
|  | L ACC |  | -3 | 53 | 2 | 4.18 |
|  | R Superior Medial Gyrus |  | 3 | 59 | 2 | 4.09 |
|  | L Superior Frontal Gyrus |  | -15 | 56 | 8 | 4.01 |
|  | L Middle Temporal Gyrus | 218 | -51 | -64 | 20 | 4.88 |
|  | L Angular Gyrus |  | -51 | -67 | 35 | 4.61 |
|  | R Precentral Gyrus | 130 | 36 | -13 | 65 | 5.06 |
|  | R Postcentral Gyrus |  | 36 | -28 | 47 | 3.97 |
|  | R Superior Temporal Gyrus | 102 | 51 | -49 | 23 | 4.46 |
|  | R Middle Temporal Gyrus |  | 57 | -52 | 20 | 4.35 |
| Advanced group |  |  |  |  |  |  |
| conflict > non-conflict | L IFG (p. Triangularis) | 482 | -39 | 20 | 26 | 5.28 |
|  | L Precentral Gyrus |  | -51 | 5 | 38 | 4.88 |
|  | L Middle Frontal Gyrus |  | -33 | 53 | 14 | 4.35 |
|  | L IFG (p. Opercularis) |  | -57 | 8 | 14 | 4.21 |
|  | L Supplementary Motor Area | 314 | -3 | 11 | 56 | 5.2 |
|  | L Superior Medial Gyrus |  | -6 | 26 | 50 | 4.78 |
|  | R Superior Medial Gyrus |  | 6 | 29 | 47 | 4.7 |
|  | L MCC |  | -6 | 23 | 38 | 4.45 |
|  | R Cerebellum (VI) | 295 | 27 | -64 | -22 | 5.67 |
|  | R Cerebellum (Crus 2) |  | 9 | -88 | -31 | 4.63 |
|  | R Cerebellum (Crus 1) |  | 15 | -76 | -28 | 4.12 |
|  | Cerebellar Vermis (6) |  | -3 | -61 | -22 | 3.63 |
|  | L Inferior Parietal Lobule | 285 | -33 | -43 | 41 | 4.46 |
|  | L Superior Parietal Lobule |  | -24 | -58 | 44 | 4.45 |
|  | L Inferior Parietal Lobule |  | -27 | -52 | 38 | 4.25 |
|  | R IFG (p. Opercularis) | 236 | 48 | 11 | 26 | 4.89 |
|  | R IFG (p. Triangularis) |  | 42 | 26 | 11 | 4.37 |
|  | R Insula Lobe |  | 33 | 29 | -1 | 4.35 |
|  | R Caudate Nucleus | 124 | 15 | 5 | 11 | 4.19 |
|  | L Middle Temporal Gyrus | 87 | -48 | -55 | -4 | 4.94 |
|  | L Middle Temporal Gyrus |  | -57 | -52 | 5 | 4.5 |
| non-conflict > conflict | - | - | - | - | - | - |
| Intermediate group | | | | | | |
| conflict > non-conflict | L Precentral Gyrus | 1021 | -51 | 8 | 38 | 7.99 |
|  | L IFG (p. Triangularis) |  | -45 | 32 | 17 | 6.71 |
|  | L IFG (p. Opercularis) |  | -57 | 11 | 8 | 4.42 |
|  | L Supplementary Motor Area | 709 | -3 | 14 | 53 | 7.89 |
|  | R MCC |  | 6 | 29 | 35 | 6.76 |
|  | L Superior Medial Gyrus |  | 0 | 38 | 44 | 5.73 |
|  | L ACC |  | -6 | 35 | 23 | 3.89 |
|  | L Inferior Parietal Lobule | 522 | -48 | -37 | 50 | 5.96 |
|  | L Superior Parietal Lobule |  | -24 | -70 | 53 | 5.41 |
|  | L Middle Occipital Gyrus |  | -27 | -61 | 38 | 5.01 |
|  | L Caudate Nucleus | 244 | -9 | 14 | -1 | 6.41 |
|  | L IFG (p. Orbitalis) | 165 | -33 | 35 | -16 | 6.78 |
|  | L Insula Lobe |  | -30 | 23 | -1 | 5.64 |
|  | R Pallidum | 158 | 12 | 5 | -1 | 5.24 |
|  | R Caudate Nucleus |  | 15 | 2 | 20 | 5.09 |
|  | R Cerebellum (VI) | 131 | 12 | -76 | -22 | 4.78 |
|  | R Cerebellum (Crus 1) |  | 12 | -85 | -28 | 4.48 |
|  | R Cerebellum (Crus 2) |  | 9 | -79 | -34 | 4.32 |
|  | L Cerebellum (Crus 2) |  | -3 | -79 | -34 | 3.75 |
|  | L Inferior Temporal Gyrus | 113 | -54 | -52 | -16 | 6.82 |
|  | R Middle Frontal Gyrus | 80 | 39 | 32 | 20 | 4.54 |
|  | R IFG (p. Triangularis) |  | 45 | 17 | 23 | 3.85 |
|  | R IFG (p. Opercularis) |  | 48 | 8 | 23 | 3.78 |
|  | R Cerebellum (Crus 2) | 70 | 33 | -76 | -49 | 5.28 |
|  | R Cerebellum (VIII) |  | 21 | -73 | -49 | 5.14 |
| non-conflict > conflict | R Superior Frontal Gyrus | 87 | 15 | 53 | 32 | 4.65 |
|  | R Superior Medial Gyrus |  | 12 | 59 | 23 | 4.37 |
| Beginner group |  |  |  |  |  |  |
| conflict > non-conflict | L Precentral Gyrus | 472 | -48 | 5 | 38 | 5.47 |
|  | L IFG (p. Opercularis) |  | -33 | 14 | 26 | 4.95 |
|  | L IFG (p. Triangularis) |  | -45 | 20 | 23 | 4.59 |
|  | L Supplementary Motor Area | 253 | -6 | 11 | 56 | 6.17 |
|  | R MCC |  | 9 | 26 | 35 | 4.13 |
|  | L ACC |  | -12 | 29 | 29 | 4.12 |
|  | R Superior Medial Gyrus |  | 9 | 26 | 44 | 4.12 |
|  | L Inferior Parietal Lobule | 107 | -42 | -43 | 47 | 4.82 |
|  | L Inferior Parietal Lobule |  | -48 | -34 | 41 | 4.51 |
|  | L Superior Parietal Lobule | 69 | -24 | -64 | 47 | 4.15 |
|  | L Superior Occipital Gyrus |  | -24 | -67 | 32 | 4.08 |
| non-conflict > conflict | - | - | - | - | - | - |

**Table S2.** The results of rhyme judgement task
