## Supplementary material for "Proficiency-Dependent Reorganisation of Language and Control Networks during Second Language Processing: An fMRI Study of Korean–English Bilinguals": Table S3 (from another version of this preprint: version 1, 2026-08-19)

**Supplementary table 3**

| contrast | Region | cluster | MNI coordinate | | | T |
| --- | --- | --- | --- | --- | --- | --- |
|  |  |  | x | y | z |  |
| All |  |  |  |  |  |  |
| conflict > non-conflict | L IFG (p. Opercularis) | 480 | -45 | 11 | 20 | 6.21 |
|  | L Middle Orbital Gyrus |  | -39 | 56 | -4 | 5.2 |
|  | L IFG (p. Triangularis) |  | -42 | 38 | 14 | 5.15 |
|  | L IFG (p. Orbitalis) |  | -45 | 47 | -13 | 4.7 |
|  | L Precentral Gyrus |  | -42 | 11 | 41 | 4.41 |
|  | L Middle Frontal Gyrus |  | -36 | 17 | 53 | 3.88 |
|  | L Inferior Parietal Lobule | 269 | -42 | -49 | 47 | 4.87 |
|  | L Inferior Parietal Lobule |  | -36 | -49 | 41 | 4.81 |
|  | L Middle Occipital Gyrus |  | -27 | -64 | 35 | 4.07 |
|  | L Supplementary Motor Area | 142 | -6 | 20 | 47 | 5.37 |
|  | L Superior Medial Gyrus |  | -3 | 35 | 35 | 4.65 |
|  | R Cerebellum (VII) | 130 | 24 | -76 | -49 | 4.94 |
|  | R Cerebellum (Crus 1) |  | 45 | -70 | -37 | 4.09 |
|  | R Cerebellum (VIII) |  | 30 | -67 | -52 | 3.78 |
|  | R Cerebellum (Crus 2) |  | 39 | -64 | -43 | 3.71 |
|  | R Cerebellum (VI) |  | 24 | -61 | -34 | 3.6 |
|  | R Inferior Parietal Lobule | 82 | 33 | -55 | 47 | 4.05 |
|  | R Inferior Parietal Lobule |  | 48 | -52 | 53 | 4.03 |
| non-conflict > conflict | L Lingual Gyrus | 258 | -9 | -79 | -4 | 4.38 |
|  | L Precuneus |  | 0 | -64 | 20 | 3.9 |
|  | L Calcarine Gyrus |  | -12 | -61 | 17 | 3.89 |
|  | L PCC |  | 0 | -49 | 26 | 3.78 |
|  | R Calcarine Gyrus |  | 24 | -58 | 11 | 3.75 |
|  | R Precuneus |  | 12 | -49 | 20 | 3.57 |
|  | R Cuneus |  | 18 | -70 | 23 | 3.5 |
|  | R Lingual Gyrus | 108 | 21 | -70 | -4 | 4.85 |
|  | R SupraMarginal Gyrus | 80 | 60 | -25 | 20 | 4.91 |
|  | R Superior Temporal Gyrus |  | 54 | -22 | 11 | 3.32 |
|  | L Superior Temporal Gyrus | 80 | -60 | -34 | 20 | 4.55 |
| Advanced group |  |  |  |  |  |  |
| conflict > non-conflict | L Inferior Parietal Lobule | 110 | -48 | -58 | 47 | 4.22 |
|  | L Inferior Parietal Lobule |  | -36 | -46 | 38 | 4.11 |
| non-conflict > conflict | - | - | - | - | - | - |
| Intermediate group |  |  |  |  |  |  |
| conflict > non-conflict | - | - | - | - | - | - |
| non-conflict > conflict | - | - | - | - | - | - |
| Beginner group |  |  |  |  |  |  |
| conflict > non-conflict | - | - | - | - | - | - |
| non-conflict > conflict | - | - | - | - | - | - |

**Table S3.** The results of spelling judgement task
