## Supplementary material for "Proficiency-Dependent Reorganisation of Language and Control Networks during Second Language Processing: An fMRI Study of Korean–English Bilinguals": Table S4 (from another version of this preprint: version 1, 2026-08-19)

**Supplementary table 4**

| contrast | Region | cluster | MNI coordinate | | | T |
| --- | --- | --- | --- | --- | --- | --- |
|  |  |  | x | y | z |  |
| Rhyme judgement task | | | | | | |
| Interaction | Cerebellar Vermis (7) | 118 | -3 | -64 | -25 | 13.09 |
|  | L Cerebellum (VI) |  | -6 | -67 | -16 | 12.96 |
|  | R Superior Medial Gyrus | 88 | 9 | 38 | 38 | 15.44 |
|  | L Superior Medial Gyrus |  | 0 | 41 | 47 | 12.6 |
|  | R Superior Medial Gyrus |  | 3 | 32 | 32 | 12.33 |
|  | L Superior Medial Gyrus |  | -9 | 35 | 41 | 11.94 |
|  | L Middle Orbital Gyrus | 83 | -27 | 38 | -16 | 22.34 |
|  | L IFG (p. Orbitalis) |  | -33 | 29 | -19 | 16.39 |
|  | L Temporal Pole |  | -39 | 14 | -25 | 14.92 |
|  | R Cerebellum (Crus 2) | 53 | 9 | -88 | -31 | 15.95 |
|  | L Cerebellum (Crus 2) |  | 0 | -79 | -34 | 12.25 |
|  | L SupraMarginal Gyrus | 50 | -54 | -49 | 26 | 14.04 |
| Spelling judgement task | |  |  |  |  |  |
| Interaction | L Cerebellum (IV-V) | 109 | -9 | -40 | -13 | 13.07 |
|  | L Cerebellum (III) |  | -6 | -37 | -16 | 12.35 |
|  | L Thalamus | 93 | -12 | -13 | -1 | 14.86 |
|  | R Middle Frontal Gyrus | 56 | 33 | 53 | 5 | 18.5 |
|  | R Superior Frontal Gyrus |  | 30 | 62 | 11 | 10.53 |

**Table S4.** The results of interaction between condition and group
